# Constructive neutral evolution explains the emergence of specialised ribosomes in diverse eukaryotes

**DOI:** 10.64898/2026.06.23.733944

**Authors:** Alan J.S. Beavan, Bulat Fatkhullin, Juan Fontana, James O. McInerney, Julie Aspden, Mary J. O’Connell

## Abstract

Throughout eukaryotic evolution, the structure of the ribosome has been highly conserved, featuring 80 common protein gene families. However, in many eukaryotes, paralogs of these proteins are present. “Specialised ribosomes” have been documented across diverse groups of eukaryotes where they play an important role in the regulation of translation of specific mRNAs. In the case of specialised ribosomes it has been documented that assembled ribosomes that contain specific paralogs can directly affect translational output. This has been proposed to contribute to the regulation of complex responses to environmental change and to coordinate cell-type specific physiology. This poses the question of whether ribosome specialisation principally emerges under an adaptive or neutral model of evolution. Using gene tree-species tree reconciliation, we test competing hypotheses regarding the evolutionary drivers of ribosome specialisation. We determine that examples of specialisation tend to emerge by independent duplication of the same ribosomal proteins in different lineages. We show that pathways to specialisation through paralog formation have arisen independent of: (i) paralog location within the 3D ribosome complex, and (ii) positive selection in these paralogs. We determine that the generalisable model of best fit for the evolution of paralog-mediated eukaryotic ribosomal specialisation is one of constructive neutral evolution. In lineages with small effective population sizes and increased complexity, the emergence and retention of ribosomal protein paralogs has provided the raw material for ratcheting and the emergence of translational regulation at the level of the ribosome.

## Introduction

Ribosomes are the macromolecular machines responsible for protein synthesis in all cellular life. Current data suggest that only one additional ribosomal protein (RP) family, the P3 protein, has joined the ribosome since the last common ancestor of eukaryotes, an addition that is confined to certain plant lineages (Gutiérrez, et al. 2004). Overall, the eukaryotic ribosome is considered to be a large and highly conserved complex composed of four or five ribosomal RNA (rRNA) molecules up to 80 RPs if we include the plant specific protein family, 79 otherwise, and not all of these RPs are incorporated into fully synthesised ribosomes (Emmott, et al. 2019). However, while the fundamental architecture of the ribosome has remained remarkably stable throughout eukaryotic evolution, its constituent proteins and rRNAs have diversified considerably across lineages – the ribosome has evolved in significant ways since eukaryogenesis including the emergence of specialisation.

Eukaryotic ribosomes are more complex than their prokaryotic counterparts in four main ways. These are: 1) the number of proteins involved in the mature ribosome complex, which is 57 in bacteria and either 79 or 80 in eukaryotes (34 of which are shared across domains) (Lecompte, et al. 2002); 2) the length and structure of rRNA molecules, which have undergone further expansion in complex lineages of eukaryotes (Hsiao, et al. 2009; Petrov, et al. 2014; Rauscher and Polacek 2024); 3) the number of associated proteins involved in ribosome biogenesis (Strunk and Karbstein 2009); and 4), the expansion in the number of paralogs of RPs (Nakao, et al. 2004). Indeed, in *Saccharomyces*, 60 of the 79 RP gene families retain duplicates derived from an ancient whole-genome duplication (Wolfe and Shields 1997) and paralogs of RPs are prevalent across eukaryotes (Nakao, et al. 2004). Why this complexity has arisen exclusively in eukaryotes has been the source of debate, given the existence of prokaryotes for substantially longer timescales and their tendency towards higher effective population sizes where smaller adaptive benefits are felt by selection (Lynch and Conery 2003). The ubiquity of ribosomes across cellular life means that adaptive explanations of eukaryotic ribosomal complexity have been proposed. Under the adaptive framework, expansions in complexity increase the ribosome’s functional capacity, for example by allowing the recruitment of context-specific translation factors, without disrupting the conserved, functional core (Fox 2010; Petrov, et al. 2015). However, an alternative explanation is constructive neutral evolution (CNE) whereby neutral or slightly deleterious mutations are allowed to drift to fixation according to the nearly neutral theory of molecular evolution (Kimura 1983; Ohta 1992). As these nearly neutral interactions become cemented through interdependency, a “complexity ratchet” occurs, wherein a reduction in complexity cannot occur without deleterious effects (Stoltzfus 1999; Gray, et al. 2010; Lynch 2024). Population genetic theory states that these mutations likely carry a small adaptive cost through higher likelihood of mutation and are more likely to be retained in lineages with low effective population sizes - characteristic of eukaryotes (Lynch and Conery 2003). As such, CNE has offered as an explanation for the complexity of the eukaryotic ribosome, enabling the increased number of proteins involved, the action of rRNA expansion segments, and the expansion of the ribosome biogenesis pathway, all compared with prokaryotic ribosomes (Gray, et al. 2010; Muñoz-Gómez, et al. 2021). Proponents of the CNE explanation of ribosomal complexity argue that there is no evidence that eukaryotic ribosomes are faster or make fewer errors than prokaryotic ribosomes but necessarily incur a higher energetic cost (Lynch 2024).

Currently missing from this theoretical work is an explanation of the duplication and retention of RPs in so many eukaryotes. The most common fate of paralogs is loss (Lynch and Conery 2000). However, when two paralogs are retained and have different sequences (as we see in the case of RP family evolution), under the framework of fitness landscapes it is possible that one of the retained paralogs is better suited to translation of certain mRNAs (Wright 1932). Multicopy gene families including RPs are generally rare in prokaryotes (Nakao, et al. 2004; Makarova, et al. 2005; see also Lilleorg, et al. 2019) therefore, duplication and retention of RP genes in eukaryotes and subsequent divergence could be understood as becoming more complex under a CNE framework just as other aspects of the ribosome can (Stoltzfus 1999).

Our understanding of eukaryotic translation is undergoing a major change. We now know that ribosomes are not uniform protein-producing machines; instead, they display compositional variation both within and among cell types. The structural diversity generated through neutral accretion may have provided the evolutionary substrate for ribosome heterogeneity, enabling the emergence of tissue-, condition-, or lineage-specific specialised ribosomes. **Ribosome specialisation** requires that compositionally distinct ribosomes, have a functional effect on the translational output, *i.e.* specialised ribosomes regulate translation (Xue and Barna 2012; Shi, et al. 2017; Guo 2018; Beavan, et al. 2025; Milenkovic and Novoa 2025; and for an alternative view point, Ferretti and Karbstein 2019).

One of the most widely studied mechanisms underlying ribosome specialisation is paralog switching, whereby duplicated RP genes are differentially incorporated into ribosomes. RP paralog switching within ribosomes can modulate the translation of defined mRNA pools, linking ribosome heterogeneity to developmental and physiological outcomes (Milenkovic and Novoa 2025). For example, gene duplication of the large-subunit RP eL22, has occurred independently in fruit flies, mammals, and yeast, and in each lineage, paralog switching and ribosome specialisation has been demonstrated with varying degrees of evidence (Komili, et al. 2007; O’Leary, et al. 2013; Mageeney and Ware 2019; Li, et al. 2022). In mice, the germ-cell-specific paralog eL39 is required for male fertility (Li, et al. 2022). Similarly, a heart- and skeletal-muscle-specific paralog uL3 directs translation of contraction-related proteins (Milenkovic, et al. 2023) which, from a structural perspective, has in turn been associated with proximity to the peptide-exit tunnel of the ribosome, implying that even subtle sequence divergence in key structural locations can yield specialised ribosome phenotypes. Similarly, paralog switching in *Drosophila* is enriched around peptide exit tunnel and near the mRNA channel, though the extent to which these changes enable specialisation remains to be determined (Hopes, et al. 2021). In *Arabidopsis*, the regulation and inclusion of different uL30 paralogs in ribosomes appears to partly coordinate the cold response (Martinez-Seidel, et al. 2021). Intriguingly, paralog switching in the same RP family across distantly related eukaryotes occurs (eL22, uL30), with paralog pairs sometimes differing by only a few amino acids yet conferring distinct translational profiles (Crawford and Pavitt 2019; Malik Ghulam, et al. 2022). The persistence of these paralogs has been attributed both to adaptive pressures – where paralog diversification supports regulatory control over translation (Dinman 2016), and to non-adaptive dynamics consistent with CNE and population-genetic theory (Force, et al. 1999). Throughout this paper, the concept of specialised ribosomes is defined as pools of ribosomes that are heterogeneous according to the paralogs they recruit and have different translational signatures.

Here, we ask whether paralog-mediated ribosome specialisation is a product of adaptive innovation (selecting for translational control), or whether it has largely emerged through constructive neutral evolution, *i.e.* the accumulation and entrenchment of neutral molecular complexity. We integrate phylogenomic, structural, and simulation-based studies, and provide an explicit test of the constructive neutral evolution model for the emergence and persistence of ribosome specialisation.

## Materials and Methods

### Gene family construction and gene tree inference

Datasets and gene families were specified according to methods outlined graphically (Figure 1) and described here. Annotated genomes were downloaded from Ensembl (Dyer, et al. 2024) and NCBI (Sayers, et al. 2025) for 356 eukaryotic species spanning all major eukaryotic groups and enriching for sampling around model organisms (principally *Arabidopsis thaliana, Homo sapiens, Mus musculus, Drosophila melanogaster, Saccharomyces cerevisiae*). To ensure maximum genome quality we used BUSCO (Manni, et al. 2021) (eukaryota_odb10), and any genomes with >30% of expected genes missing were excluded. A set of exceptions to the missing gene filter were included either because their missing genes can be explained by their unusual life histories or because they are the most complete genome available for a given clade. This accounts for many parasite genomes where we see high rates of gene loss and “missing genes” are expected (Khalaf, et al. 2024). In total, we retained 243 genomes (**Supplementary Table S1**), the “**243_taxon_set**”. The corresponding species tree was generated according to previous scientific studies (**Supplementary information**). In addition, to avoid erroneously inferring gene duplication events due to potentially mis-specified species tree topology, we excluded species if they contributed a species tree node that is contentious in the literature. We extracted a subset of 83 species from the 243 species dataset that contained no controversial relationships and prioritised the presence of model organisms. This 83-species set featured only opisthokonts (animals and fungi) and archaeplastida (plants and green algae) between which the root was placed.

**Figure 1:**
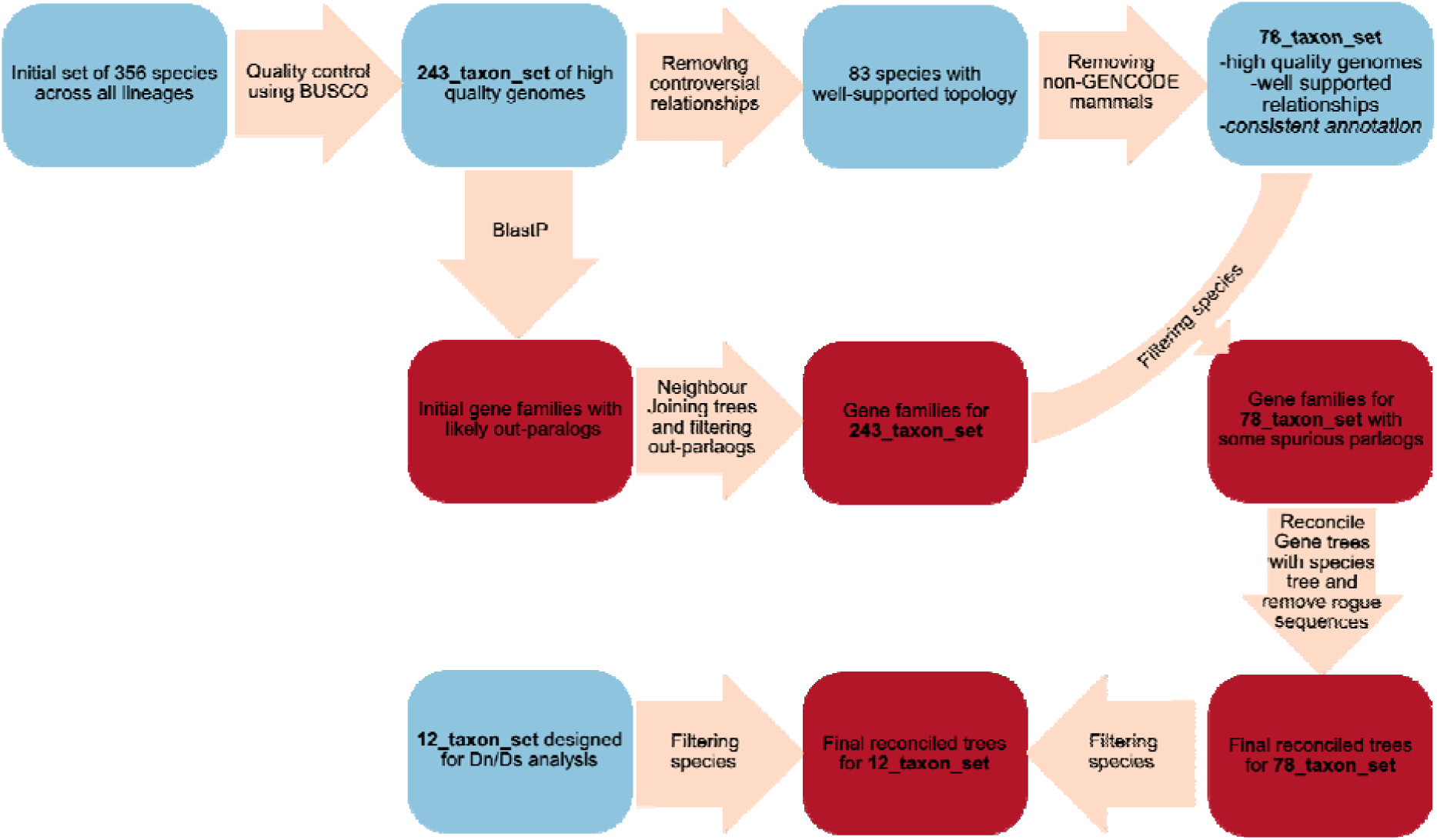
Data flow, taxon sampling and gene tree inference implemented in this study. Each step of quality control is indicated by arrows linking sets of taxa (blue boxes) or gene families (red boxes). Technical aspects of each step are indicated in the main text.

A final filtering step was applied due to differences in how mammal genomes were annotated. The origin of mammals is characterised by many retrotransposition events which resulted in protein coding genes being copied back into the genome from the processed transcript, resulting in single exon genes with homology to the gene from which it was retrotransposed (Zhang, et al. 2003). However, different annotation pipelines treat these retrotransposed genes differently, with the GenCode annotations removing them and others maintaining them as “genes” (Mudge, et al. 2024). The result of this is that most mammal genes have an abnormally high average copy number among animals, but mouse and human do not (**Supplementary Figure S1**). Therefore, to avoid inferring infeasible levels of duplication and loss in the mammal lineage, all non-GenCode annotated mammal genomes were removed. The resulting 78 taxon tree was retained for subsequent gene tree-species tree reconciliation and subsequent analysis and is referred to as the **78_taxon_set** throughout.

RP sequences, referred to as “seeds” here, were downloaded for all eukaryotic species in the Ribosomal Protein Gene Database (Nakao, et al. 2004). Each seed sequence, or set of seed sequences, was compared to each of the genomes in the 243_taxon_set using BLAST (BLASTP) (Altschul, et al. 1990; Altschul, et al. 1997) with e-value set to 10-3. As some of the seed sequences may belong to the same gene family, blast results were merged where families overlapped in gene content by ≥50%. Multiple sequence alignments were made for gene families using MAFFT version7.310 (Katoh and Standley 2013) with default settings, and a neighbour joining tree was constructed for each family using FastTree version 2.1.11 (Price, et al. 2010). Trees were rooted at the midpoint. As an initial filtering step, highly distant paralogs identified from these neighbour joining trees were removed to maintain more computationally manageable datasets. Sequences from the clade representing the most recent common ancestor of each seed sequence were taken forward. If this set of sequences was less than 2000, the set of sequences was expanded to include the smallest clade over 2000 sequences from this tree. This strategy allows for the inclusion of out-paralogs but, more importantly, ensures that all the members of the orthologous gene family are included. Subsequently, the longest canonical peptide sequence for each gene was selected by comparing the lengths of all proteins produced from the same gene.

To avoid inferring erroneous gene duplication events due to uncertainty in the species tree, all subsequent analyses were performed on the gene trees from the 78_taxon_set. The selected sequences for each gene family were re-aligned using MAFFT with default settings and trimmed using trimAL 1.4 (Capella-Gutierrez, et al. 2009), removing sites with gaps in at least 50% of the sequences. This was preferred to the automated methods strict and strictplus, which produced alignments that were too short to infer evolutionary relationships, and gappyout because the alignment length was longer in most cases when removing sites with 50% gaps without substantially changing the alignment quality (**Supplementary Figure S2**). A model of protein evolution was inferred on a gene family basis using IQTree 2.1.2 and ModelFinder (Kalyaanamoorthy, et al. 2017; Minh, et al. 2020). IQTree generates a neighbour joining tree as part of this process. Each RP alignment was then used as input to GeneRax 2.1.3 (Morel, et al. 2020). GeneRax also uses a species tree to infer a joint likelihood of the evolution of the protein sequences under the model and the relationships of the proteins given the species tree. The reconciliation model within GeneRax was Duplication and Loss. Transfer was forbidden because, while rare instances of rDNA horizontal gene transfer have been documented in some eukaryote lineages (Yabuki, et al. 2014), none have for RPs and it is unlikely that transfer of these genes would occur over the large evolutionary distances covered in this set of taxa. Each gene family used the substitution model inferred by ModelFinder and a starting gene tree as inferred by IQTree using neighbour joining methods.

Finally, gene trees were pruned according to **two** criteria:

**(1) Paralogous genes were removed by retaining only species that descend from a common orthologous ancestor**. For RP genes, where a range of eukaryotic species were used as seed sequences, the common ancestor of all genes with sequence identical to at least one of these seed sequences was treated as the orthologous gene family common ancestor.
**(2) Genes present in a topology that are hard to explain under a model of gene duplication and loss only.** This refers to instances where the inferred phylogenetic position of a given gene requires an ancient duplication and subsequent loss in many lineages and retention in their own. This causes some duplication events to be erroneously inferred as old. Accordingly, software was developed (https://github.com/alanbeavan/rogue_gene_removal/releases/tag/v1.0) to infer the number of loss events needed to explain the presence of each gene in the gene tree under a model of duplication and loss. While the inclusion of most genes can be explained by no extra loss events, there are some which require a substantial number of loss events. For this reason, genes that require an extra 6 loss events, a threshold inferred from the distribution of extra loss events needed for a gene’s inclusion (**Supplementary Figure S3**), which resulted in 489 genes out of 115,071 being removed from further analysis. We acknowledge that these genes may be interesting, emerging from: A) horizontally transferred from a distant lineage, B) sequence contaminants in the sequencing of one of the genomes sampled in this study, or C) out-paralogs retained through previous pruning steps. However subsequent analyses are more robust with their removal. Following these two rounds of pruning, the gene trees were re-reconciled with the species tree using GeneRax (--strategy EVAL).

### Characterising specialised ribosomes via paralog switching

We have extracted known cases of specialisation from the literature; this set contains 12 pairs of RP paralogs across 5 gene families (see **Table 1** and references therein). These cases were further split into two sets; a **strict** set where the evidence is strongest and a **relaxed** set containing the **strict** set alongside cases where the evidence is less strong. The **relaxed** set includes genes where changes in RP paralog abundance are correlated with changes in protein expression, but heterogeneous populations of ribosomes have not been directly implicated. For all pairs of paralogs implicated in specialisation (**strict** or **relaxed**), the branch of the species tree on which the paralogs diverged was inferred by finding the common ancestor of all the species that retain at least one copy of the pair of paralogs. Further, to assess our hypothesis that ribosome specialisation is facilitated by shifts in selective pressure of paralogous RP sequences, we performed selective pressure analyses on a subset of species in all RPs. To do this comparison we generated a subset of the alignment that contained only model organisms with some additional species to reduce the length of long internal branches (*Danio rerio, Mus musculus, Homo sapiens, Caenorhabditis elegans, Tribolium castaneum, Drosophila melanogaster, Drosophila simulans, Saccharomyces cerevisiae, Saccharomyces kudriavzevii, Arabidopsis lyrata, Arabidopsis thaliana, Chlamydomonas reinhardtii*). This process yielded subsamples containing 12 species, referred to hereafter as the **12_Taxon_Set** (see **Supplementary Table S1** for a full set of genomes and the categories to which they belong).

**Table 1:** The set of known cases of paralog-mediated specialised ribosomes used in this study.

| Gene family | Species | Set | Explanation |
| --- | --- | --- | --- |
| eL22/RPL22 | *D. melanogaster* | **strict** | *In vivo* evidence that eL22 and eL22-like have specialised, non-interchangeable functions, as they associate with distinct subsets of mRNAs and drive selective translation programs during *Drosophila* spermatogenesis, with eL22-like unable to fully compensate for loss of eL22 (Mageeney and Ware 2019). |
| eL39/RPL39 | *M. musculus* | **strict** | The testis-specific eL39 paralog, eL39L, is required for spermatogenesis and forms part of a testis-specific ribosome with distinct structural and functional properties that selectively regulates germ-cell protein synthesis and cannot be functionally replaced by canonical eL39-containing ribosomes (Zou and Qi 2021; Li, et al. 2022). |
| uL3/RPL3 | *M. musculus* | **strict** | Though discrepancies in findings remain (Grimes, et al. 2023; Milenkovic, et al. 2023), uL3L is a striated muscle-specific RP paralog that forms functionally distinct ribosomes with altered translation elongation dynamics and transcript-specific effects on cardiac protein synthesis (Chaillou, et al. 2016; Shiraishi, et al. 2023). |
| uL30/RPL7 | *S. cerevisiae* | **strict** | uL30/RPL7 paralogs' products are differentially acetylated with the hyper-acetylated form on exposure to a drug, shifting translation from short to long ORFs which may promote tolerance to the drug (Malik Ghulam, et al. 2022) . |
| eL22/RPL22 | *S. cerevisiae* | **relaxed** | ASH1 mRNA translation depends specifically on eL22A rather than eL22B, indicating functional specialization of eL22 paralogs, although direct evidence for distinct mRNA-selective ribosome populations was not provided (Komili, et al. 2007). Subsequently paralog-specific regulatory and transcriptomic effects were demonstrated, including functions of eL22B that could not be compensated by eL22A (Abrahámová, et al. 2018), providing additional support for the idea that the two eL22 paralogs contribute differently to gene expression and potentially to ribosome specialization. |
| eL22/RPL22 | *M. musculus* | **relaxed** | eL22 regulates expression and ribosomal incorporation of eL22L1, providing evidence that eL22 paralogs contribute to ribosome heterogeneity in mammals (O'Leary, et al. 2013). |
|  |  |  | Dynamic replacement of eL22 by eL22L1 during spermatogenesis forms part of a sperm-specific ribosome program required for specialised germ-cell functions (Li, et al. 2022). |
| uL30/RPL7 | *A. thaliana* | **relaxed** | Paralog-specific shifts in uL30 gene expression are associated with cold response. These changes in paralog expression are incorporated into translationally active ribosomes. The authors propose that these cold-induced paralog exchanges could remodel the ribosomal polypeptide exit tunnel and thereby constrain translation elongation of specific mRNAs, although direct demonstration of uL30-specific translational selectivity is not provided (Martinez-Seidel, et al. 2021). |
| uL16/RPL10 | *M. musculus* | **relaxed** | uL16L is a component of heterogeneous germ-cell ribosomes that cannot be functionally replaced by canonical ribosomes and is essential for male fertility. uL16's role in specialisation is less well characterised than in eL39 but mirrors its patterns in expression (Tu, et al. 2020; Li, et al. 2022). |

For each gene family tested, the amino acid sequences of the genes present in the small subset of species were aligned using MAFFT (Katoh and Standley 2013) with default settings. Subsequently, the nucleotide coding sequences were superimposed onto this amino acid alignment using PAL2NAL V.14 (Suyama, et al. 2006), generating a codon alignment. Using codon-based alignments and gene family trees, BUSTED version 3.0 (part of the HyPhy suite of tools (Murrell, et al. 2015)) was used to assess whether the evolution of the nucleotide sequences was significantly better explained by allowing some nucleotide sites to evolve with a ratio of non-synonymous substitutions per nonsynonymous site to synonymous substitutions per synonymous site (Dn/Ds) > 1 on a selection of several branches. These branches were defined as those immediately following a duplication event that had led to retention of both paralogs in at least one of the descendent species (see **Supplementary Figure S4**). For every duplication event, each paralogous lineage was selected as the “foreground” separately and together, meaning three hypotheses were put forward for each duplication event:

1. One branch emerging from the duplication
2. The alternative branch emerging from the duplication
3. Both branches together.

The P-value was extracted from BUSTED’s output files for each case.

In addition to this, all alignments were re-coded according to the Dayhoff-6 scheme (Dayhoff 1972). This means that each amino acid was converted to a symbol representing a set of amino acids that behave similarly, but nucleotide (codon) alignments remained unchanged. This was done because the timescales covered in this study are large and without recoding, there is a risk that nucleotide substitutions would be saturated, reducing the power of conventional Dn/Ds analyses (Susko and Roger 2007). By implementing BUSTED on the re-coded amino acid alignments, we asked whether the evolution of these emergent paralogs was characterised by an increase in the ratio of Dayhoff-recoded non-synonymous changes to Dayhoff-recoded synonymous changes. To illustrate this, change of TCT to ACT is a non-synonymous change converting a serine to a threonine. However, as serine and threonine belong to the same Dayhoff-6 group (Embley, et al. 2003), this is a synonymous Dayhoff-recoded change.

### 3D structural analysis

To assess whether rates of duplication were enriched by the peptide exit tunnel and the mRNA channel, ribosome structures from representative model species, representing the highest-resolution and most complete models available for each species at the time of analysis were downloaded from the Protein Data Bank (Zardecki, et al. 2022) for Human (pdb_00006y57 (Bhaskar, et al. 2020)), yeast (pdb_00008cgn (Milicevic, et al. 2024)), and *Arabidopsis* (pdb_00004v7e (Gogala, et al. 2014)), and the distance from each protein to each landmark (exit tunnel or mRNA channel) was taken as the minimum distance between two residues (either amino acid or nucleotide) in each molecule. For RPs, the locations of the residues were defined by their alpha carbon atoms. Likewise, the peptide exit tunnel was defined by the alpha carbon atoms in a nascent peptide. For the mRNA channel, the space was defined by the phosphorus atoms in the bound mRNA molecule. To assess the relationship between duplication rate and distance from each of these coordinates a Pearsons’ correlation test was performed. Structures were visualised using ChimeraX-1.6.1 (Meng, et al. 2023).

## Results

### Extensive duplication, divergence and loss in the evolution of eukaryotic ribosomal proteins

To determine the duplication and loss rates of RP genes per species per branch on the **78_taxon_set** (see materials and methods) (**Figure 1**), we reconciled the inferred gene trees with the species tree of known topology and calculated a duplication rate of 0.26 per branch and loss rate of 0.25 per branch (**Supplementary Table S3**). By reconstructing the most parsimonious number of copies of each RP coding gene, we determined that, in terms of protein content, the ribosome was already a complex and sophisticated machine in LECA, featuring 79 of the 80 genes examined (the exception being the plant-specific large-subunit protein P3 which is most likely to have emerged in the ancestor of land plants). This approach also allowed us to estimate that of the 79 RPs present, 70 were present in single copy and 9 contained two or more paralogs in LECA (*i.e.* a ratio of single copy to two or more paralogs of 70:9). Applying a similar approach for ancestral gene copy number state reconstructions we asked similar questions for all other major clades (**Figure 2**). We determined that the ratios of single copy to two or more paralogs of RPs in the most recent common ancestor (MRCA) of animals, fungi and plants (Streptophyta) were 54:25, 66:13, and 56:23 respectively. High numbers of paralogs in *Arabidopsis* and *Saccharomyces* (**Figure 2**) are likely due to retention of paralogs from independent ancestral whole genome duplication (WGD) events specific to those lineages (Wolfe and Shields 1997; Clark and Donoghue 2018) (indicated on **Figure 2A**). We also show disparity in rates of duplication across gene families (**Figure 2B**) and lineages (**Figure 2C**), with plants generally having the most duplication events inferred across gene families and fungi the least. However, this pattern is not repeated across all gene families, with some animal RP families experiencing more duplications in animals than in plants and some featuring more duplications in fungi than in animals (**Figure 2D**). Overall, these patterns show that while the RP repertoire was established in LECA, its subsequent evolutionary history has been shaped by gene and lineage-specific duplication dynamics rather than a uniform pattern.

**Figure 2:**
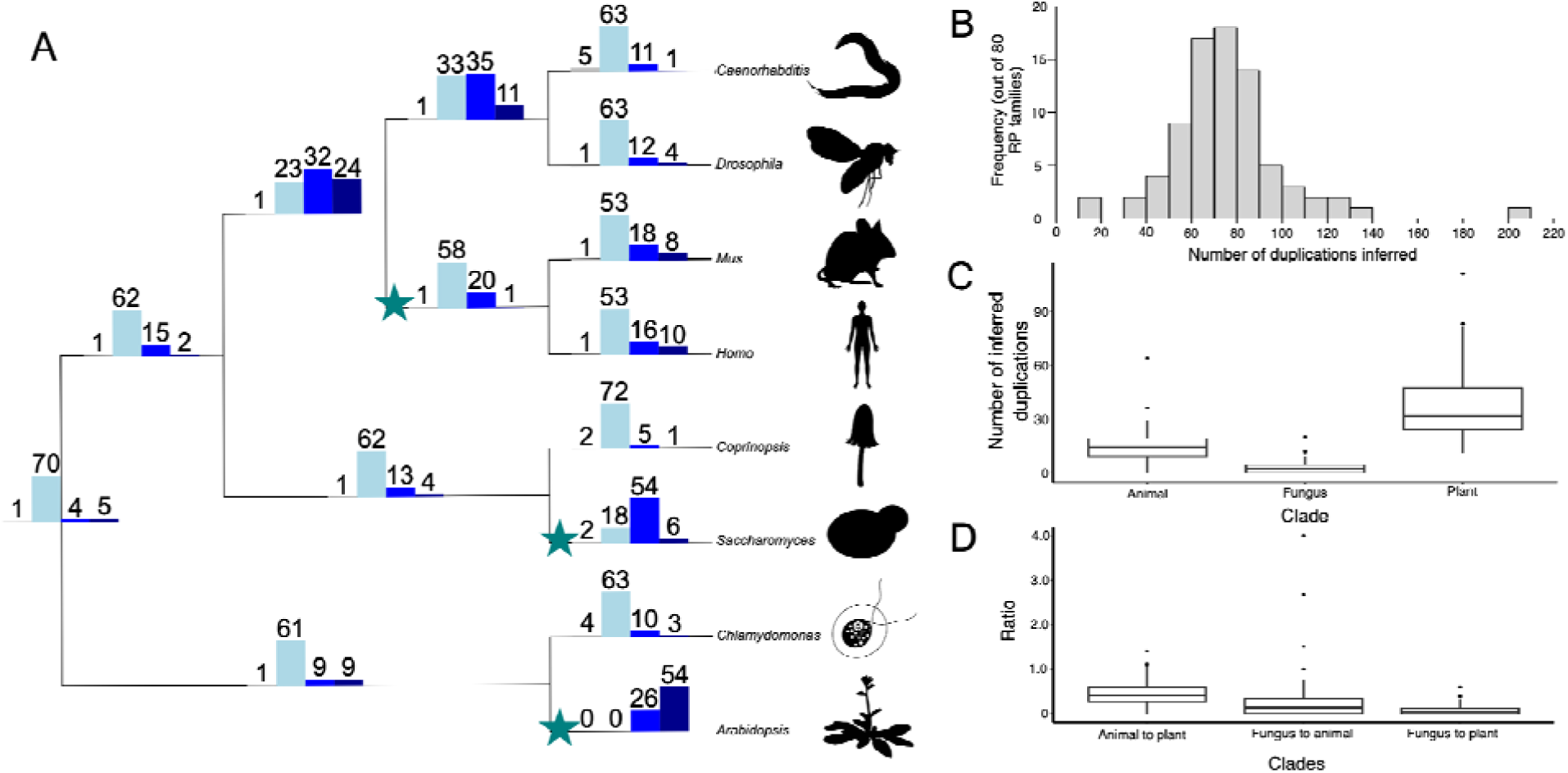
The profile of RP gene singletons and paralogs depicted on the cladogram of eight genera subsampled from the full set of 78 species. **(A)** The histogram on each branch indicates the number of RP genes (out of eighty) on that branch. Each histogram contains four bars of distinct colour, from left to right across each histogram these represent the number of RP genes (i) absent (present in zero copies; grey), (ii) present in single copy (light blue), (iii) present in two copies (blue), and (iv) present in more than two copies (dark blue). Lineages featuring at least one well supported WGD event are indicated by a star. **(B)** Histogram of the number of duplications inferred for each RP gene family. **(C)** The number of duplications inferred in all RP families across three major eukaryote clades. For each clade, boxes represent the interquartile range, the thick solid line the median number of inferred duplication and the whiskers extend to the furthest datapoints up to 1.5 times the interquartile range. **(D)** The ratios of duplications inferred per RP gene family between three major eukaryote clades with boxes and whiskers specified as in (**C**).

### Rates of RP gene duplication are not associated with functional centres of the ribosome

Further to heterogeneity in gene duplication and loss events across lineages, we asked whether duplication rates were enriched in regions of the ribosome to which gene products are mapped.

There is also heterogeneity across gene families, with each RP family experiencing duplication between 14 and 202 times across eukaryotic history (**Supplementary Table S3**). The number of duplications across families (excluding the plant specific P stalk protein RPLP3) ranged from 0 to 64 (31 species) in animals, from 0 to 20 (12 species) in fungi, and from 11 to 112 (30 species) in plants. To determine whether this variation in duplication history was associated with the position of the RP within the 3-D structure of the eukaryotic ribosomal complex we parsed the gene trees reconciled with the species tree and inferred the number of duplication events for each gene in each of the three major clades, Animals, Fungi and Plants. The resulting profiles of duplication were mapped to a clade-typical 3-D ribosome structure (**Figure 3**).

**Figure 3:**
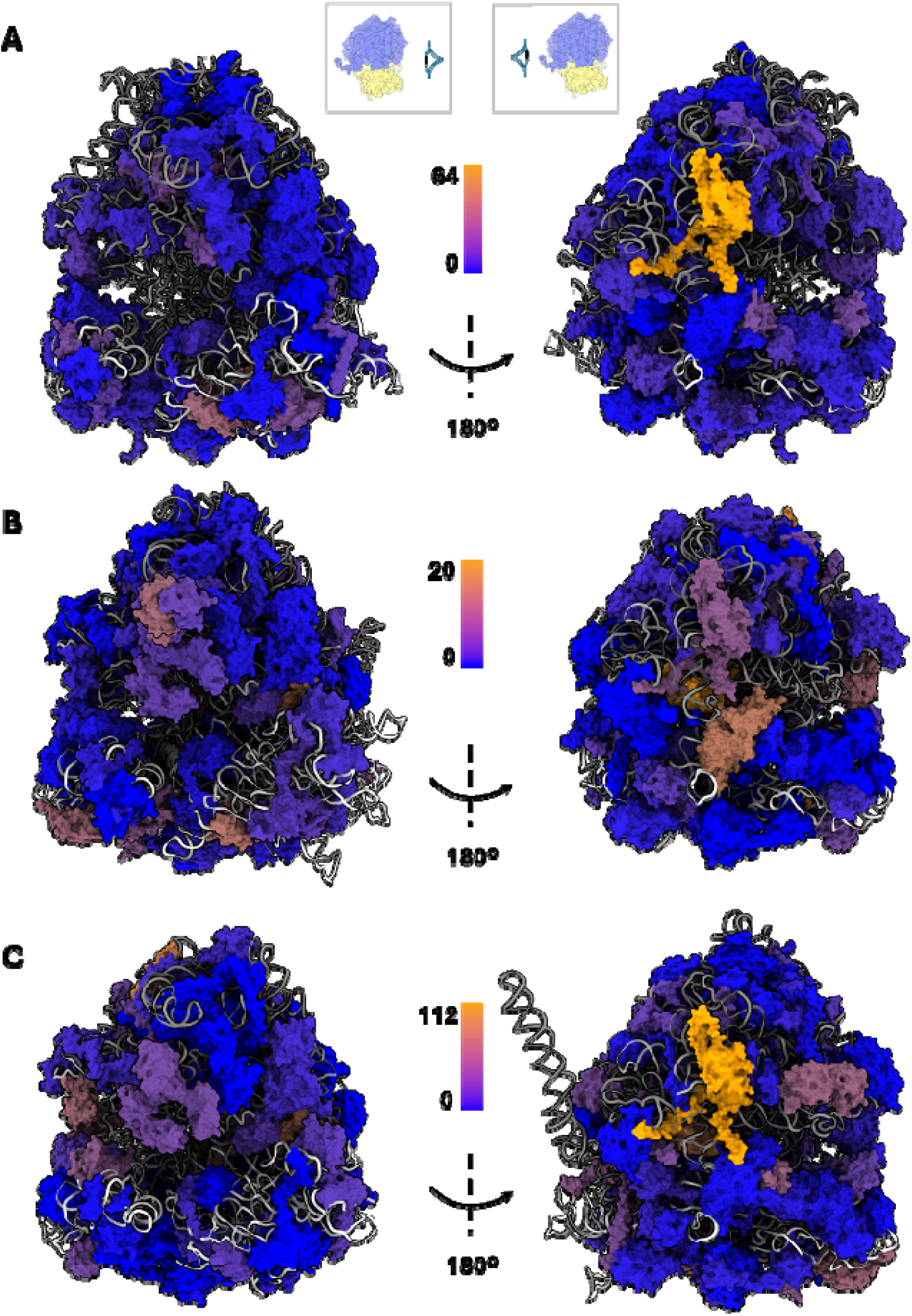
Three-dimensional ribosomal structures displaying duplication profile of RPs. The constituent RPs of the ribosomal complex are represented as colour surfaces, where the colour of each protein depicts the number of duplications inferred see corresponding scale per panel. Low levels of gene duplication in blue to high levels of gene duplication in orange. **(A)** animals (PDB pdb_00006y57), **(B)** fungi (PDB pdb_00008cgn) and **(C)** plants (PDB pdb_00004v7e). The RNAs are represented as cartoon; 18S rRNA is coloured white; 5S, 5.8S, 28S, tRNAs and mRNAs are coloured in different shades of grey in each structure.

Given that specialisation has been proposed to be associated with RPs in specific functional centres of the ribosomal complex - the mRNA channel and the nascent peptide exit tunnel (Hopes, et al. 2021)- and given that RP gene duplication can lead to paralog switching that can also lead to specialisation, we tested whether RP gene duplication events were enriched in spatial regions of the ribosome. Finding an enrichment of duplication events near functional centres might suggest RP gene duplication and specialisation are intrinsically linked. To test this hypothesis, the minimum distance between each protein and each functional centre (exit tunnel and mRNA channel) was compared with the number of duplications inferred for that protein in the clade (animal, fungi or plant) from which the 3D ribosome structure was derived. For each clade, and for each functional centre, the two variables were compared using Pearson’s correlation, revealing no significant correlation between duplication rate and distance from functional centres (P>>0.05 before correcting for multiple tests using False Discovery Rate). This result was replicated when the average distance (across all amino acids per protein) was compared with the number of duplications inferred instead of the minimum distance (Pearson’s correlation P >> 0.05). This suggests that either A) specialisation is not necessarily more likely to occur in proximity to the mRNA channel or peptide exit tunnel, or B) gene duplication is not closely correlated with specialisation, that is, there are other factors governing RP gene duplicability. Our findings suggest no pattern of duplication correlating with spatial regions and specialisation across the plant, animal and fungal ribosome structures.

### Paralogs involved in specialised ribosomes emerge convergently in diverse lineages and over vast evolutionary timescales

Of the five gene families suggested in the **relaxed** set (including examples where evidence for ribosome specialisation is strongest and examples where the evidence is less strong), proposed to have been implicated in specialised ribosomes by paralog switching, three of them have been characterised by multiple duplication events in the lineages found to be characterised by specialisation (**Table 1**). In total, we identify 12 duplication events with retained paralogs shown to be involved in specialisation. These duplication events range from relatively recent lineages, such as eL22 in the ancestor of *Drosophila*, through to the duplication of *Saccharomyces* uL30 in an ancestor of all extant fungi. In both eL22 and uL30, where specialisation has been suggested in multiple lineages, we find that these families duplicated independently in each lineage in which specialisation has been detailed (**Figure 4**).

**Figure 4:**
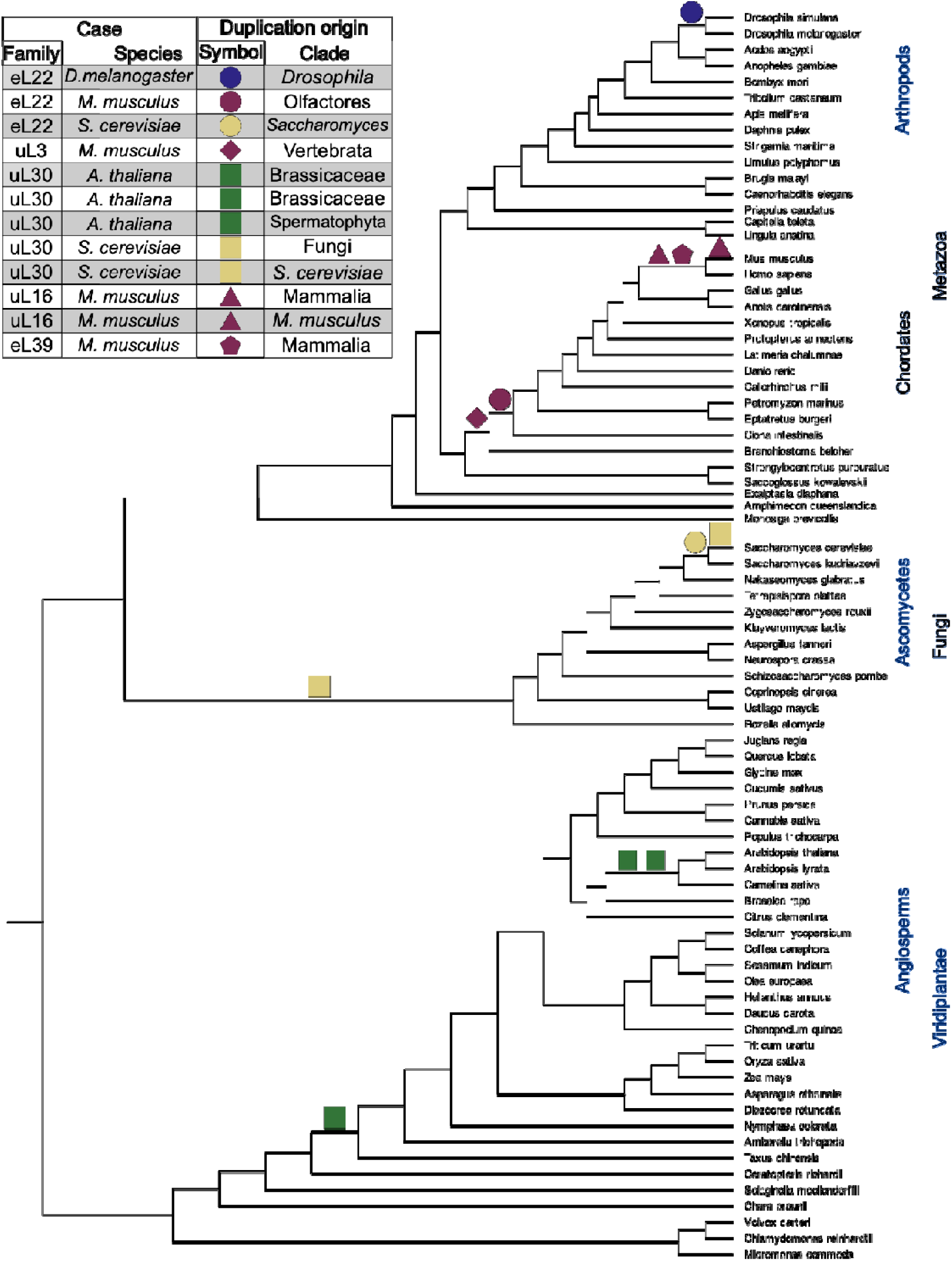
The duplication of genes in known cases of specialisation. The species tree of the **78_taxon_set** is shown with major clades indicated. Known cases of ribosome specialisation via paralog duplication from the **relaxed** set are represented in the table, imposed over the species tree branch on which they have been inferred to have duplicated.

### Paralogs involved in specialised ribosomes do not have unique patterns of adaptation

To test whether positive selection could be driving the emergence of specialised ribosomes by paralog switching with neofunctionalisation, we assessed selective pressure variation in paralogs of RP genes where both copies were retained in at least one of a set of 18 species. We asked whether these duplication events resulted in an increase in the classic measurement of selective pressure variation, *i.e.* is the evolution of these emergent paralogs better explained by at some sites that evolve with Dn/Ds > 1 (evidence of positive selection) compared with the evolution of the gene family as a whole. Retained paralogs were split into those where there is already evidence that the paralog is suggested to be implicated in specialisation (“known”, *i.e.* 12 gene duplication events across 5 gene families (**Figure 4**)) and those where no evidence has been documented (“unknown”, *i.e.* 395 gene duplication events across all RP gene families). Here, the “known” set is equivalent to the **relaxed** set (**Table 1**). Across both “known” and “unknown” subsets of the data, and across the single branches and pairs of branches tested (as described previously), we find variation in Dn/Ds across the dataset along with evidence of positive selection, *i.e.* the model for the foreground is a significantly better fit when a Dn/Ds > 1 is permitted (**Figure 5c&d – positive selection? = Yes, and Supplementary Table S3**). To investigate these patterns in more detail, we compare the “known” RP paralogs directly with the “unknowns” and observe that the proportion of single branch tests indicating positive selection is not significantly different in the “known” and “unknown” cases (Fisher’s Exact Test – Odds Ratio = 1.40, P = 0.48) (**Figure 5a&c**). This is also true when we test both paralogs together (**Figure 5b&d**) (Fisher’s Exact Test – Odds Ratio = 1.40, P = 0.65). We also asked whether re-coding the amino acids according to Dayhoff-6 categories changed this result. As above, known cases were no more likely to be better explained by allowing a ratio of Dayhoff-recoded non-synonymous substitutions than Dayhoff-recoded synonymous substitutions > 1 than unknown cases (one branch – Fisher’s Exact Test – Odds Ratio = 0, P = 1.0; both branches – Fisher’s Exact Test – Odds Ratio – 0.46, P = 0.7; **Supplementary Figure S5**).

**Figure 5:**
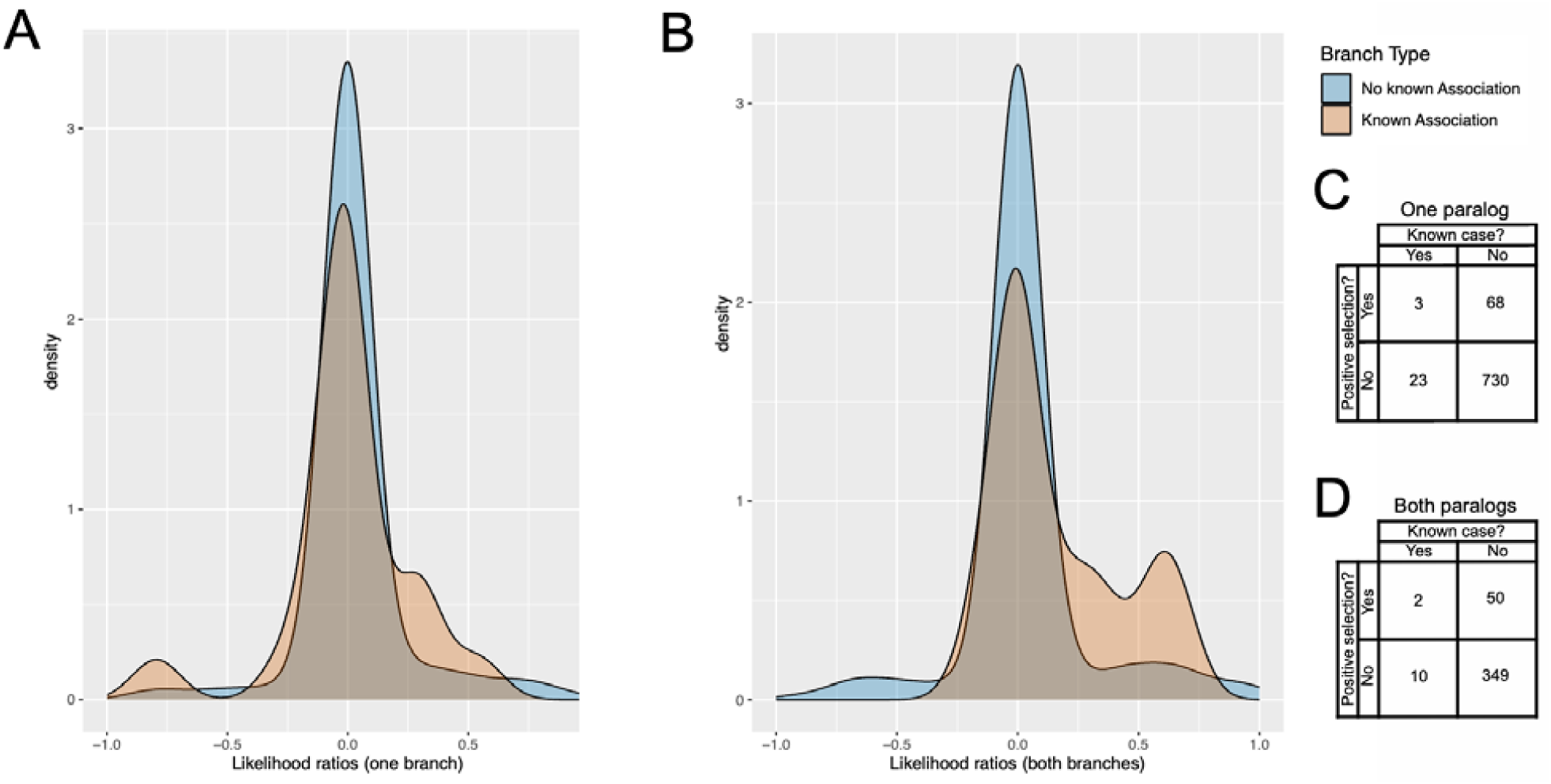
Likelihood of branches being under positive selection in RP paralogs known to be associated with specialisation as compared to those of unknown status. **(A)** and **(B)** represent the likelihood ratios returned from the positive selection analysis, using BUSTED. Instances where there is no known association between the paralogs and specialised ribosomes are shown in blue, while instance where a known association has been documented are shown in orange (**Table 1**). In **(A)** we are testing one branch emerging from duplication as the foreground and allowing Dn/Dn. > 1 in the alternative model. The ratios plotted are between the likelihood of the alternative model to the null model so tests where the likelihood ratio is greater than 0 mean that the alternative model has a higher likelihood (though this does not mean the relationship is statistically significant). In **(B)** both branches emerging from duplication form the foreground. **(C)** Contingency table showing how many duplication events are significantly better explained by allowing Dn/Ds > 1 in one of the emergent paralogs in cases associated with specialisation and cases not known to be associated with specialisation. **(D)** Contingency table showing how man duplication events are statistically significantly associated with positive selection when both paralog immediately following duplication were considered the foreground.

To determine if this result is robust to our definition of a “known case” we carried out the following analyses: In addition to the **relaxed** set (**Table 1**), we tested the **strict** set, with stronger evidence for involvement in specialised ribosomes. Due to low numbers of **strict** “known” cases (4), a power analysis was performed. Given the support for enhanced Dn/Ds in the “unknown cases”, a Wilcoxon test would require three out of four of the “both branches” cases to have been associated with adaptive evolution and three out of the eight cases for the “one branch” tests. However, neither the “one branch” nor the “both branches” tests were associated with a signature of positive selection (allowing some sites to evolve with Dn/Ds > 1) (Fisher’s Exact Test – Odds Ratio = 0, P = 1.0 and Odds Ratio = 0, P = 1.0 respectively) (**Supplementary Figure S6**). Overall, these results support a scenario where RP paralogs known to be involved in specialisation are no more likely to have signatures of positive selection than other RP paralogs and that this result is robust to our definition of a “known” case.

## Discussion

In this study, we ask whether the emergence and retention of RP paralogs, and their subsequent contribution to specialised ribosomes, is driven by adaptive evolution, or has arisen via constructive neutral evolution. We have shown that the evolution of the protein component of eukaryotic ribosomes is characterised by heterogeneous rates of gene duplication across gene families (**Supplementary Table S3**), across lineages (**Figure 2, Supplementary Table S3**), and across the 3D structure of the ribosome (**Figure 3**), with no obvious association between pattern of duplication and function. The vast majority (1,055 out of 1233) of tests for selective pressure variation revealed RP paralogs are not associated with higher rates of adaptive evolution immediately following their duplication as determined by Dn/Ds analyses (**Figure 5).**

Given the absence of evidence for shifts in selective pressure, we propose that this complexity (*i.e.* ribosomal specialisation) has likely emerged under a nearly neutral process by either of the following two scenarios: (i) genetic drift has led to the retention of both copies, or (ii), that whole genome duplication events have meant that pairs of RP paralogs were immediately fixed in the population and not lost later, possibly being retained due to constraints imposed by stoichiometric ratios of RPs and rRNAs (Makino and McLysaght 2010). The WGD explanation may work for lineages such as vertebrates (Ohno 1970; Dehal and Boore 2005), *Saccharomyces* (Wolfe and Shields 1997), most plants (Clark and Donoghue 2018), but does not hold for other lineage such as *Drosophila*, where no such WGD process has occurred.

The importance of ribosomal function means that RPs unsurprisingly display strong signatures of purifying selection (where natural selection purges mutations). Our observations do not suggest adaptive neofunctionalisation driving the emergence of specialised ribosomes in diverse eukaryotic lineages and are instead consistent with the protein component of the ribosome becoming more complex in terms of the paralogs present, through a CNE process. These results complement similar observations in rRNA expansion segments, which are repetitive sections of rRNA molecules that tend to be more repeated in more complex forms of life (Petrov, et al. 2014). The results are also in agreement with population-genetic theory predictions that gene duplicates are more likely to be retained in lineages with low effective population sizes, where purifying selection is inefficient and slightly deleterious mutations can drift to fixation (Lynch and Conery 2003; Innan and Kondrashov 2010). Indeed, multicopy gene families are far more prevalent in eukaryotes than in prokaryotes, which typically experience larger effective population sizes and more efficient purifying selection (Makarova, et al. 2005). Our observations of RP paralog retention in eukaryotes are consistent with a view of gene duplication that does not rely on selection to fix paralogs in the first place. We do not propose a mechanism of RP gene duplication and retention here but note that previous studies have highlighted roles for WGD, retrotransposition, and a preference for the immediate retention of duplicates to buffer against haploinsufficient, dominant mutations in diploids, all of which are compatible with the results of this study (Zhang, et al. 2003; Kondrashov and Koonin 2004; Mullis, et al. 2020). In the case of WGD, genes in dosage balance, such as those coding products that act in complexes, including ribosomes, are more likely to be retained after WGD to maintain chemical stoichiometry (Makino and McLysaght 2010). This suggests WGD is a potentially strong driver in the retention of RP paralogs and fitting our data showing high paralog counts in WGD associated lineages (**Figure 2**). It is notable that in comparison to prokaryotes where there is some involvement of RPs in paralogous pairs (*e.g.* bL31 and bL36) that may be important translational regulators (Lilleorg, et al. 2019), eukaryotes retain much higher proportions of multi-copy gene families including RPs. This suggests that our findings may not be limited to eukaryotes despite their effective population size differences. Future testing of prokaryotes would augment the findings presented in this study.

If the emergence and retention of RP paralogs is non-adaptive, how then do specialised ribosomes emerge, and how are they retained by evolution? Paralog-mediated specialised ribosomes (*i.e.* those pools of ribosomes that are heterogeneous according to the paralogs they recruit and have different translational signatures) occur in a diverse set of model organisms including *Drosophilia melanogaster, Mus musculus, Saccharomyces cerevisiae* and possibly *Arabidopsis thaliana* (**Table 1**) and recent evidence has shown that this functional heterogeneity is important in coordinating complex functions (Beavan, et al. 2025). Indeed, whilst specialised ribosomes are not restricted to eukaryotes, this study and others show that the paralogs that facilitate specialisation have lineage-specific, independent origins (Hopes, et al. 2021). For example, eL22 has been implicated in specialisation in *Drosophila* (Mageeney and Ware 2019), *Saccharomyces* (Komili, et al. 2007), and mammals (O’Leary, et al. 2013). Likewise, uL30 paralogs have been implicated in specialised ribosomes in yeast and plants (Martinez-Seidel, et al. 2021; Malik Ghulam, et al. 2022). Our results suggest these paralogs (i.e. eL22 and uL30) emerged independently (**Figure 4**). However, complex structures driven by neutral evolutionary forces are not precluded from being co-opted for an adaptive benefit once established (Muñoz-Gómez, et al. 2021), and we do not rule out the role of adaptive evolution in promoting and maintaining specialised ribosomes after the paralogs were established. Indeed, it is clear than once established, RP paralogs have become an important means by which translational regulation through ribosome specialisation arises, *e.g.* expressing the wrong uL30 paralog in muscle can significantly reduce myotube growth in mice (Chaillou, et al. 2016). However, our findings show that the immediate diversification of paralog pairs post duplication is not associated with an increase in adaptive evolution.

We find diffuse duplication rates of RPs are not associated with the emergence of specialised ribosome associated paralogs (**Supplementary Table S3**) and there is no correlation between the proximity of RP paralogs to either the peptide exit tunnel or the mRNA channel (which have been previously suggested as important coordinates for the emergence of specialised ribosomes (Hopes, et al. 2021)). Together these findings mean that the duplication process itself is not being driven by the adaptive benefit of specialised ribosomes. Instead, it appears that RP paralogs emerge across the 3D structure of the ribosome and only some of those duplicates have the capacity to later bestow the property of specialisation on the organisms in which they are found. In other words, duplication is not targeted and specialisation is emergent from available variation. The presence of a diversity of RP paralogs in eukaryotic genomes increases the evolutionary potential of the eukaryotic ribosome, allowing certain lineages to repeatedly co-opt specific paralogs for specialised translational functions (Pigliucci, 2008). Therefore, the emergence of specialised ribosomes through RP gene duplication represents an interesting case of CNE, one where genetic drift has independently driven similar sets of paralogs across vast evolutionary distances that have been co-opted for a translational regulation function that is now important for normal organismal function (Chaillou, et al. 2016; Hopes, et al. 2021; Martinez-Seidel, et al. 2021; Norris, et al. 2021; Beavan, et al. 2025).

Protein coding evolution is not the only way that paralog functions can diverge and, indeed, given previous gene expression work on RP paralogs, we would expect that the regulation of these paralogs has diverged in interesting and significant ways, *e.g.* in mammalian uL3 one paralog is expressed at greater levels in heart and skeletal muscle (Milenkovic, et al. 2023). However, the divergence of expression patterns in sets of paralogs, or regulatory subfunctionalisation (Lynch and Conery 2000) alone cannot explain how these they are able to change the translation signature of a specialised ribosome, which must also require functional divergence.

An appreciation that ribosomes are actively regulating translation has only recently emerged and there are many potential discoveries for which this study will be relevant, such as new instances of paralog switching (Milenkovic and Novoa 2025), medical implications of heterogeneous ribosomes (Rivalta, et al. 2025), and a deeper understanding of how transient protein factors may enable specialised ribosome function (Cates, et al. 2025). Our study represents a framework in which to view other complex functions such as ribosome biogenesis. Prokaryotes do not require non-ribosomal proteins *in vitro,* but some assembly factors are present *in vivo* (Shajani, et al. 2011; Birikmen, et al. 2021), in direct comparison to eukaryotic ribosome biogenesis which typically requires at least 200 extra-ribosomal proteins (Thomson, et al. 2013). Future studies may ask whether such complexity emerged under a similar CNE (or non-adaptive) process. We acknowledge that instances of ribosome specialisation by paralog switching remain undiscovered across eukaryotes. This study will aid the investigation of new cases of paralog-mediated specialisation and provides a framework for understanding their evolution.

In summary, specialised ribosomes most likely emerged under a non-adaptive model. The widespread duplication and retention of RP paralogs across eukaryotes is consistent with a general model in which low effective population sizes permit the fixation of nearly neutral changes, leading to the accumulation of ribosomal complexity. Our results suggest that this complexity is not merely incidental but provides the substrate from which functional specialisation can arise, placing the ribosome as a tractable model system for studying CNE, and linking population-genetic processes to the emergence of translational regulatory function at the molecular and cellular level.

## Supporting information

All alignments and trees

## Acknowledgements

The authors would like to thank BBSRC sLoLa ‘RiboCode’ grant BB/X003086/1 for funding this work and all members of the RiboCode team for helpful discussions and input throughout the development of this work. We would like to extend particular thanks to Dr Anton Calabrese who provided helpful feedback in the development of this article. Work was performed on the HPC facilities at the Universities of Nottingham and Manchester. MJO’C would like to thank the Leverhulme Trust for her personal fellowship (RF-2024-492).

## Contributions

MJO’C and AJSB conceived of the study and wrote the manuscript. AJSB performed the molecular evolutionary analyses with input from MJO’C and JOMcI. JF, BF and AJSB carried out the structural analyses. All authors contributed to editing the manuscript.

## Data availability statement

Software developed in this study is available at https://github.com/alanbeavan/rogue_gene_removal/releases/tag/v1.0. All gene trees, alignments, genome accessions, duplication and loss rates across the phylogeny are supplied as supplementary files.

## Notes

### Competing Interest Statement

The authors have declared no competing interest.

