## Supplementary information for "Constructive neutral evolution explains the emergence of specialised ribosomes in diverse eukaryotes"

Supplementary information - species tree topology with reference list

Phase 2: Obtaining a species phylogeny of eukaryotes

A cladogram representing the relationships between the species included in this study was assembled according to previous studies. The relationships between major groups were constructed according to Strassert et al. 2021. (1). Nodes in our tree that were not represented in that study were resolved according to more focussed phylogenetics studies (2-36). In cases of ambiguity, we chose to use one of the accepted topologies. In cases of ambiguity, the species tree was later pruned to remove such cases of ambiguity (see Materials & Methods, main text).

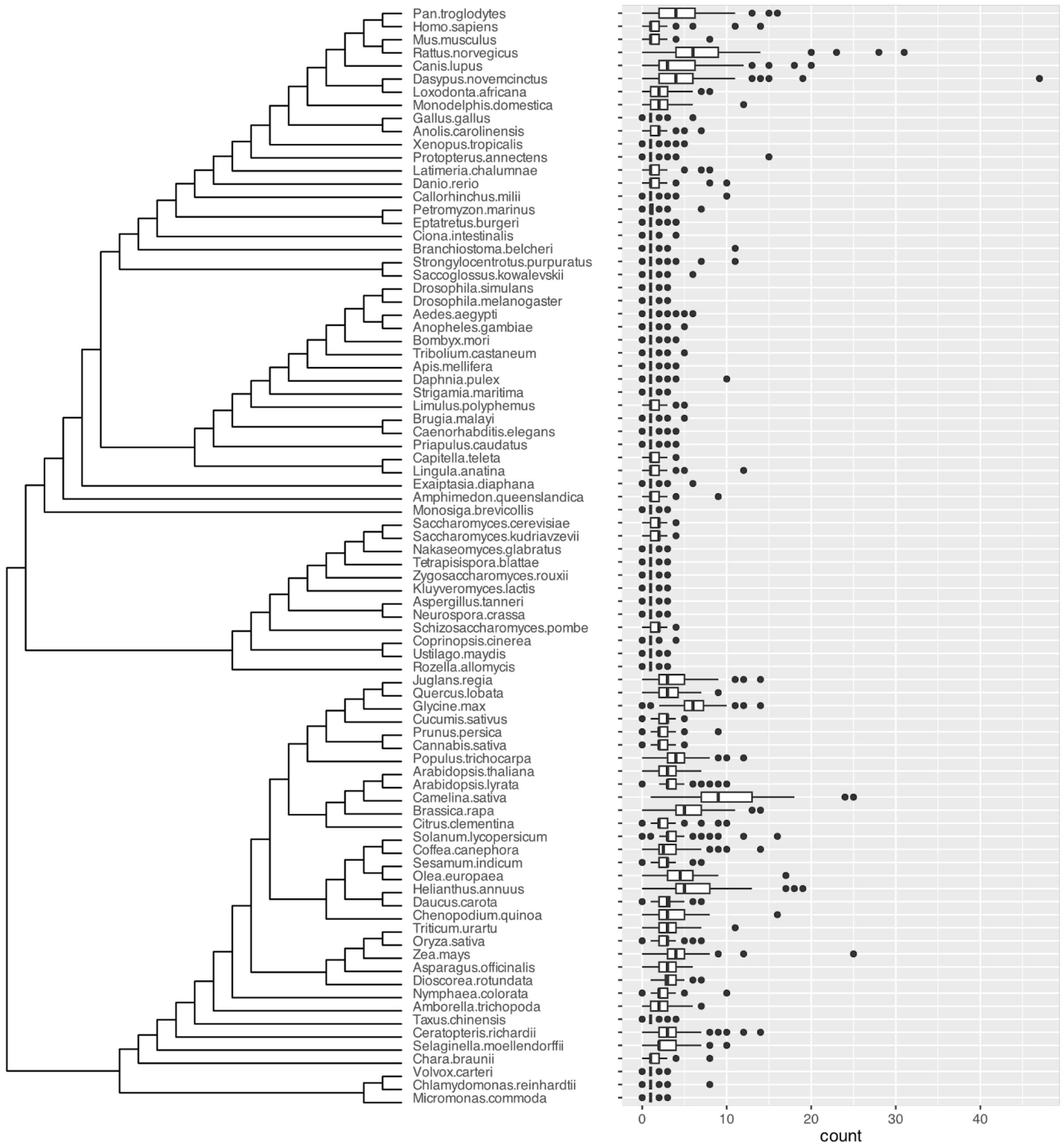


***Supplementary Figure S1:*** *The distribution of copy numbers of RP genes across the 83 taxa set defined in this study. The phylogeny on the left hand side features species corresponding to the box plots on the right hand side featuring the distribution of copy numbers over the 80 RP genes. In each box and whiskers is specified as elsewhere in this report (e.g. Supplementary Figure 2) with the number of copies represented on the X axis.*

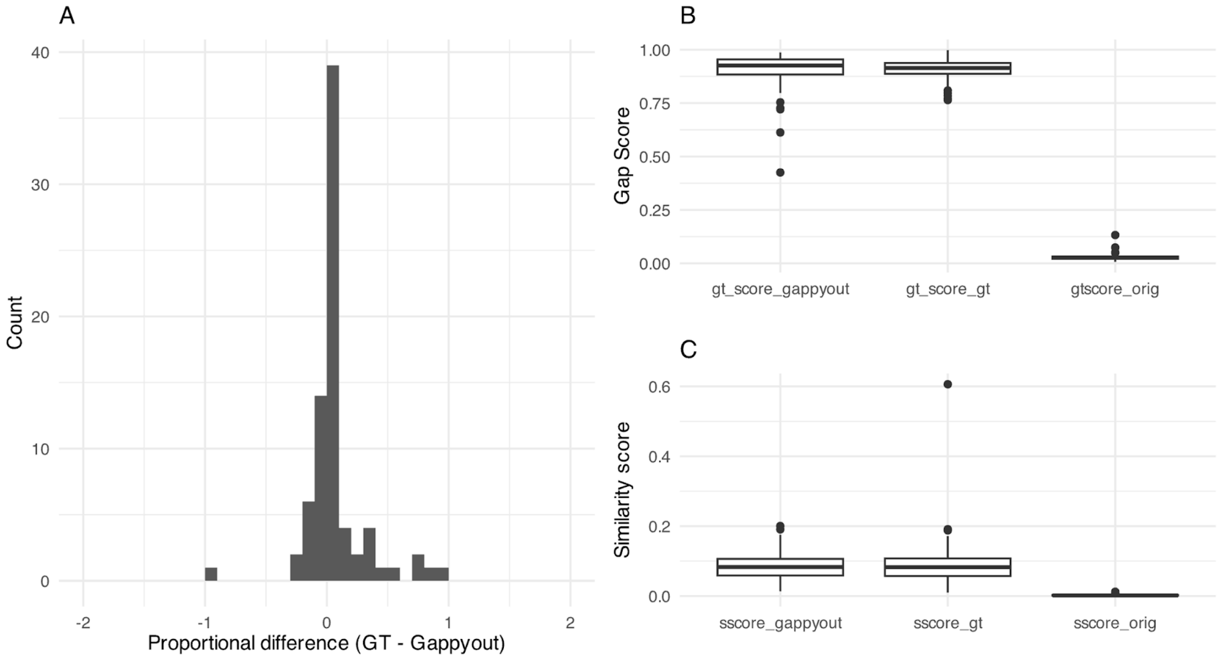


***Supplementary Figure S2: Trimming strategies and their impact on alignment length, gap proportion and similarity scores for the* 243_taxon_set.** *(A) Histogram indicating the number of RP families with the ratio of the difference between the gt50 length and the gappyout length as a proportion of the length of the shortest treated alignment out of the two strategies i.e. those > 0 are longer when treated with gt50 than gappyout. (B) The distributions of gap proportions across the original alignments, the gt50 trimmed alignments and the gappyout trimmed alignments. Each box and whiskers represent the interquartile range full range up to 1.5 times the interquartile range respectively. Points outside that range are represented as datapoints falling outside the whiskers. The median is marked with a solid line. (C) Distributions of similarity scores across the original alignment, gt50 treated alignments and gappyout treated alignments. Each box and whisker is drawn according to the same parameters as in (B).*

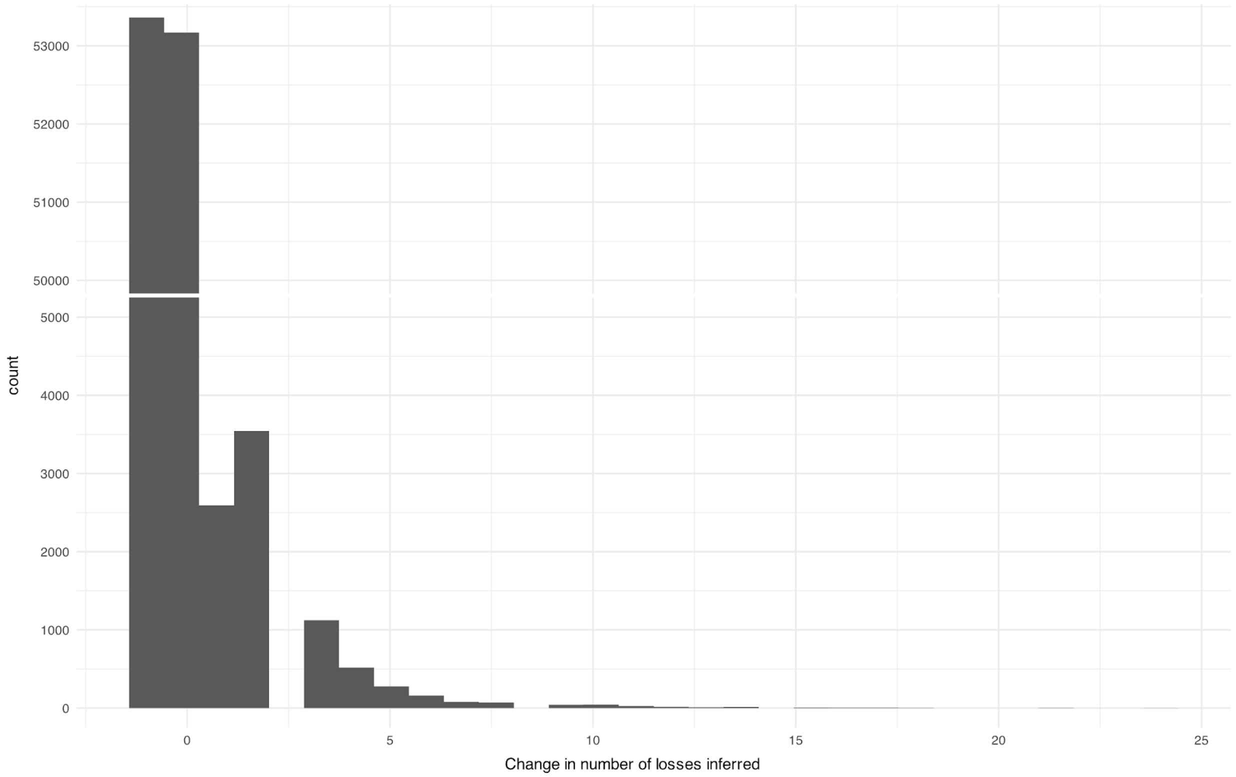


***Supplementary Figure S3: Number of losses needed for a gene's inclusion****. The plot is a histogram displaying the distribution of the difference in gene loss events needed to infer and reconcile the gene tree without a gene compared with the number inferred with it included. That is, each in the distribution was estimated by removing an individual RP gene from the alignment, inferring a gene tree and reconciling with the species tree, and extracting the difference in loss events inferred in the analysis with the gene missing and the number inferred with the full gene family. The y axis is broken between 5,000 and 50,000 where no bars terminate.*

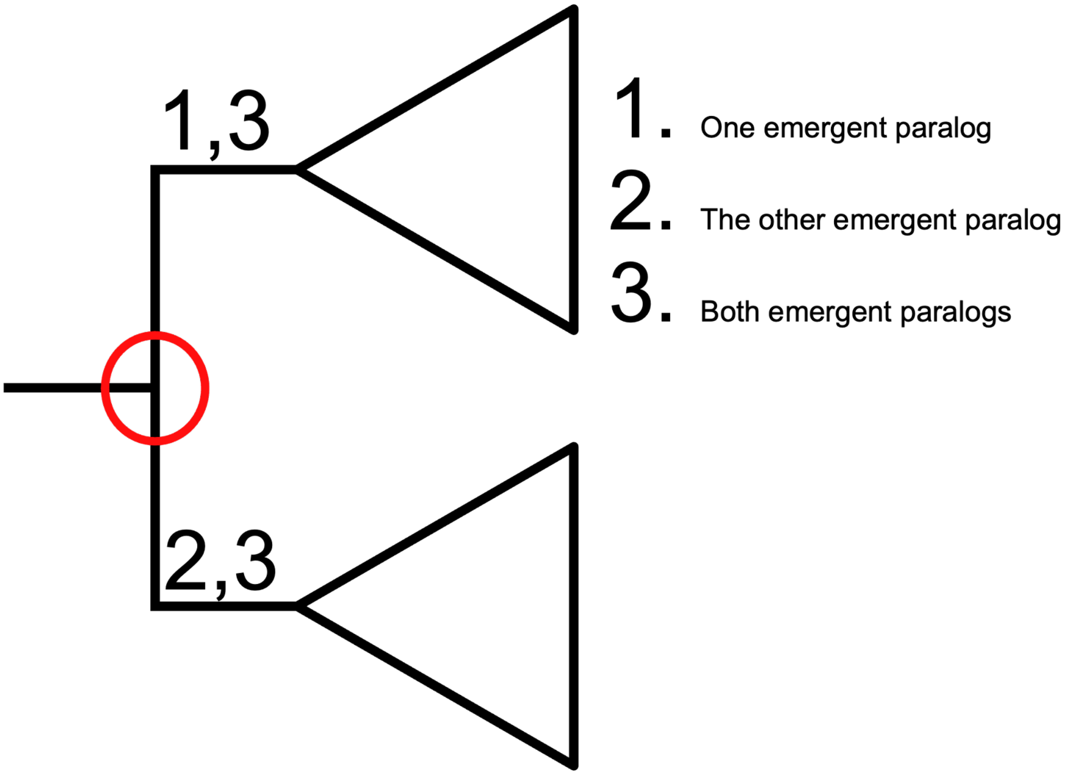


***Supplementary Figure S4: Cartoon depiction of Dn/Ds analyses performed.*** *The tree depicted represents a single ancestral gene duplicating at the node indicated with a red circle then later radiating through speciation and/or duplication as represented by the triangles. Each branch emerging after duplication is labelled with analyses 1, 2, and 3, as indicated in the main text, according to which of them is considered the foreground for this analysis.*

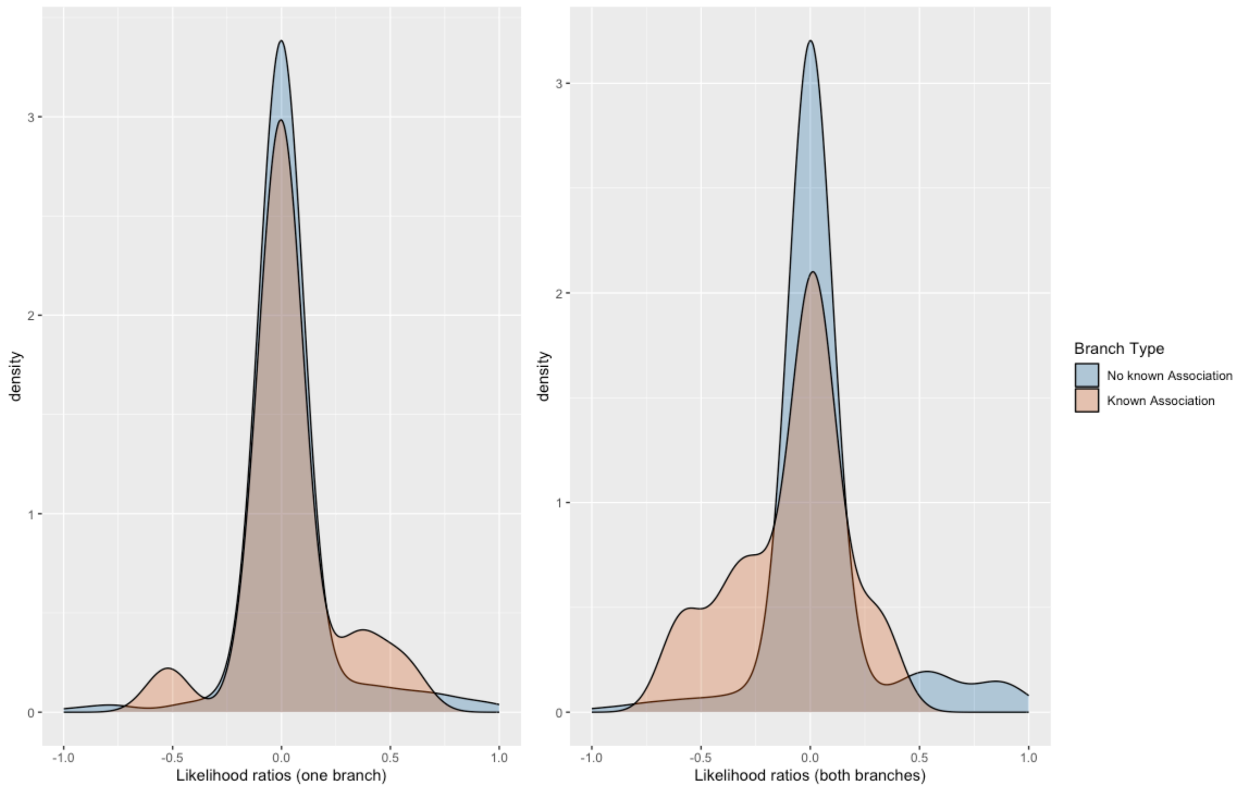


***Supplementary Figure S5: Dayhoff Category analysis for known and unknown cases of specialisation.*** *Likelihood ratios of Dayhoff-nonsynonymous to Dayhoff-synonymous substitution ratios in cases known to be involved in specialisation (orange) and not known (blue).*

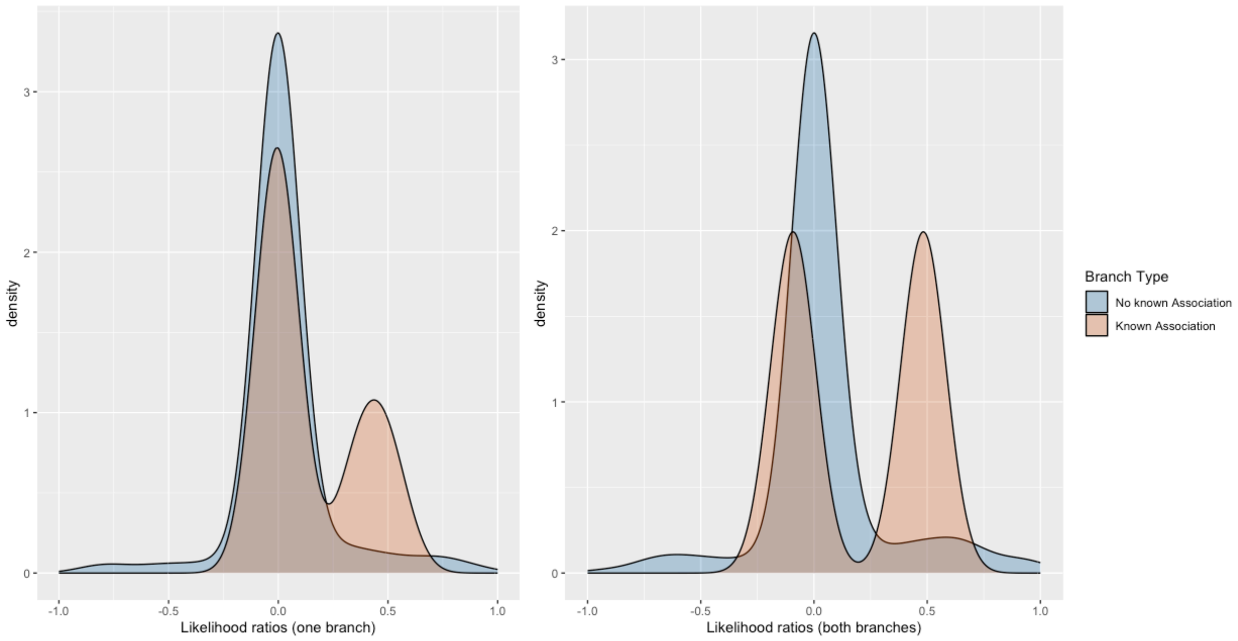


***Supplementary Figure S6: Comparison of LRTs in tests of selection for "strict" cases.*** *Likelihood ratios of nonsynonymous to synonymous substitution ratios in our* ***strict*** *cases known to be involved in specialisation (orange) and not known or known only in the* ***relaxed*** *set (blue).*

***Supplementary Table S1: Genomes used in this study, the taxon-sets they belong to, and their BUSCO completeness scores.***

***Supplementary Table S2: Duplication and loss rates per lineage.*** *Non-terminal lineages are referred to by two arbitrarily chosen tips of which that lineage represents the most recent common ancestor.*

***Supplementary Table S3: BUSTED results***. *The results of all tests of positive selection performed in this study are presented, indicated by the gene family they belong to and the set of descendants from each tip. The hypothesis is listed according to whether the left side, right side, or both sides of the duplication were specified the foreground.*
